# Memo-Patho: Bridging Local-Global Transmembrane Protein Contexts with Contrastive Pretraining for Alignment-Free Pathogenicity Prediction

**DOI:** 10.1101/2025.05.18.654712

**Authors:** Yihang Bao, Zhe Liu, Fangyi Zhao, Wenhao Li, Hui Jin, Guan Ning Lin

## Abstract

Understanding the pathogenic impact of protein mutations remains a fundamental challenge in genomic medicine, particularly for transmembrane proteins (TMPs), a functionally critical yet structurally under-annotated class that includes numerous drug targets. Current variant effect predictors often fall short in this domain due to computational burdens, reliance on multiple sequence alignments (MSAs), and limited ability to generalize to TMP-specific constraints. Here we present Memo-Patho, an alignment-free deep learning framework specifically optimized for TMP mutation pathogenicity prediction. Memo-Patho integrates global and local representations from protein language models (PLMs) and predicted structural features. Furthermore, it introduces a contrastive pre-training strategy that learns discriminative features by comparing benign and pathogenic mutations within the same protein backbone. This design enables the model to capture functional disruptions in challenging contexts without structural or evolutionary alignments. Memo-Patho outperforms state-of-the-art predictors on two types of TMP benchmark datasets (accuracy up to 0.93) and unseen-protein generalization tasks, showing strong agreement with evolutionary conservation profiles. Its predictions are further validated on a manually curated KCNQ1 dataset of 55 ion channel variants, achieving an accuracy of 0.84 and outperforming existing tools. The model also supports high-throughput scanning, efficiently analyzing over 15000 variants in under 80 minutes. Our results establish Memo-Patho as a fast, generalizable, and biologically grounded approach for variant effect prediction in TMPs. Its alignment-free design, contrastive learning paradigm, and clinical robustness mark a step forward towards scalable interpretation of TMP variation for both basic research and precision medicine applications.

## Main

The functional repertoire of a cell is executed by its proteins, whose specific roles are determined by their unique amino acid sequences and resulting three-dimensional structures [1]. Variations within these protein sequences, frequently involving the substitution of just a single amino acid residue, are common across the human population. However, even such seemingly minor alterations can profoundly disrupt protein behavior. The classic example is the Glu6Val substitution in hemoglobin’s beta chain, which leads to sickle cell anemia through altered protein solubility and aggregation [2]. Functionally compromised proteins resulting from such variants are the root cause of thousands of cataloged Mendelian diseases [3], including cystic fibrosis arising from specific CFTR protein defects (like the F508del variant) [4]. Considering the vast unexplored mutational space [5], predicting the functional impact and disease relevance of any specific amino acid change remains a fundamental challenge in molecular biology and is crucial for advancing precision medicine.

Transmembrane proteins (TMPs), comprising ∼20-30% of the human proteome [6], are integral membrane components mediating vital functions like molecular transport and signal transduction within the unique lipid bilayer environment. Consequently, single amino acid variants disrupting TMP function are implicated in numerous human pathologies. For instance, altered ion channel activity causes channelopathies affecting cardiac and neurological systems [7]. TMPs represent a large fraction of known disease genes and are the targets for over 50% of modern pharmaceuticals, highlighting the critical need to accurately assess their variant pathogenicity for diagnostics and therapeutic development [8]. However, TMPs present significant experimental hurdles: their inherent hydrophobicity complicates expression, purification, structural determination, and functional analysis compared to soluble proteins [9]. This relative scarcity of experimental data underscores the urgent need for robust computational methods specifically tailored to predict variant effects in this vital, yet experimentally challenging, protein class.

Computational prediction of variant effects has markedly evolved. Initial tools like SIFT [10] and PROVEAN [11] primarily utilized evolutionary conservation, offering foundational insights but were limited by feature scope. Subsequently, many tools began to incorporate additional features. For example, SNPs&GO introduced Gene Ontology annotations [12, 13], and SuSPect incorporated protein-protein interaction (PPI) networks [14]. STRUM [15] and FoldX [16] use protein stability to approximate the impact of amino acid variations. With advances in algorithms, machine learning based methods improved predictions by integrating diverse features using classifiers such as SVM [17], random forests [18–20], naive bayes [21, 22], and gradient tree boosting [23]. This approach leveraged broader biological information but relied heavily on feature engineering. In recent years, deep learning models have become dominant in this field. These methods can automatically learn complex representations. Typical examples include MutPred2 [24], developed by combining probability models. E-SNPs&GO [25], which utilizes language model embeddings. And AlphaMissense [26], which uses AlphaFold2 structural features [27]. There are also some deep learning tools specifically developed for TMPs, such as MutTMPPredictor [28], mCSM-membrane [29], and Pred-MutHTP [30].

However, many contemporary deep learning predictors still rely on generating multiple sequence alignments (MSAs) for evolutionary context or require protein structural data. These dependencies create bottlenecks due to computational cost or data scarcity, especially for challenging classes like TMPs. Alternatively, Protein Language Models (PLMs), leveraging large-scale self-supervised pretraining on massive sequence databases, can learn rich representations encoding structural and functional insights directly from single sequences. PLMs based methods have proven effective in diverse bioinformatics tasks. For example, SignalP 6.0, by leveraging the PLM ProtTrans [31], was the first to achieve comprehensive prediction across five types of signal peptides[32]. Additionally, Hie et al. utilized PLMs such as ESM-1b [33] to guide the evolution and variant screening of antibodies and other proteins, without requiring specific structural or functional supervision[34]. This ability to derive potent features solely from PLMs as a promising avenue for developing more scalable, generalizable, and alignment-free pathogenicity prediction tools, overcoming prior methods’ limitations.

While the impact of PLMs is increasingly evident across various bioinformatics problems, their contribution to the critical task of variant pathogenicity prediction is still developing. E-SNP&GO [25] integrated features derived from ESM-1b and ProtTrans with Gene Ontology functional annotations, employing an SVM to predict the disease association of mutations. TransEFVP [35] concatenated embeddings from four different PLMs to perform classification of mutation pathogenicity. While these tools have demonstrated some success, they often suffer from limitations, such as the direct concatenation of features into a single vector for training, which neglects local structural features. Furthermore, mutation prediction tools specifically developed for TMPs have largely yet to enter the era of PLMs.

This study investigates methods to optimize the utilization of PLM embeddings and integrate supplementary features for predicting the pathogenicity of mutations. We developed and trained Memo-Patho, a novel tool, using a dataset specifically focused on TMP mutations. Key advancements incorporated into Memo-Patho include: (1) Elimination of the need for MSAs in generating input features, which substantially improves computational efficiency and enables large-scale preliminary screening across extensive mutational landscapes. (2) Drawing inspiration from contextual understanding in natural language processing [36], PLM embeddings are leveraged distinctly as global (contextual) and local (residue-specific) representations. These are fused using a multi-head Transformer mechanism, enhancing the model’s ability to assess mutational effects within their pertinent environmental context. (3) Implementation of a contrastive learning pre-training phase prior to the final binary classification, enabling the model to more effectively capture the underlying principles governing diverse mutational impacts within a single protein sequence. (4) Integration of predicted structural features, offering insights into how mutations alter the local structural milieu of TMPs and their interactions with the surrounding environment. Rigorous validation from multiple viewpoints confirms that Memo-Patho attains state-of-the-art predictive accuracy on both internal and external benchmarks, establishing it as a valuable resource for the analysis and high-throughput screening of pathogenic mutations in TMPs.

## Results

### The Memo-Patho framework

Memo-Patho is a deep learning architecture designed for the alignment-free prediction of pathogenicity associated with mutations in TMPs. **Figure 1** shows the model structure. Its core principle involves integrating the localized impact of a mutation with its broader implications within the global protein structure and function. The architecture utilizes a dual-stream feature extraction process: (1) The Global Feature stream leverages PLMs to derive sequence embeddings, which are subsequently pooled to summarize global sequence information. This is combined with predicted structural features to form a comprehensive global feature vector representing the overall protein context; (2) Concurrently, the Local Feature stream focuses on the mutation site, extracting sequence and structural features specifically at this locus using PLMs and structure predictors. A dedicated Local-Global Feature Fusion Module then synergistically processes these features. Both local and global feature vectors undergo flattening and linear transformation. Positional encoding is added to retain sequential or spatial information before they are fed into a Transformer block. This block, via its multi-head self-attention mechanism [37], dynamically weighs feature elements and models complex interdependencies between the local mutation site and the global protein context. The output from the Transformer is passed through a final projection layer, yielding a unified representation.

**Figure 1:**
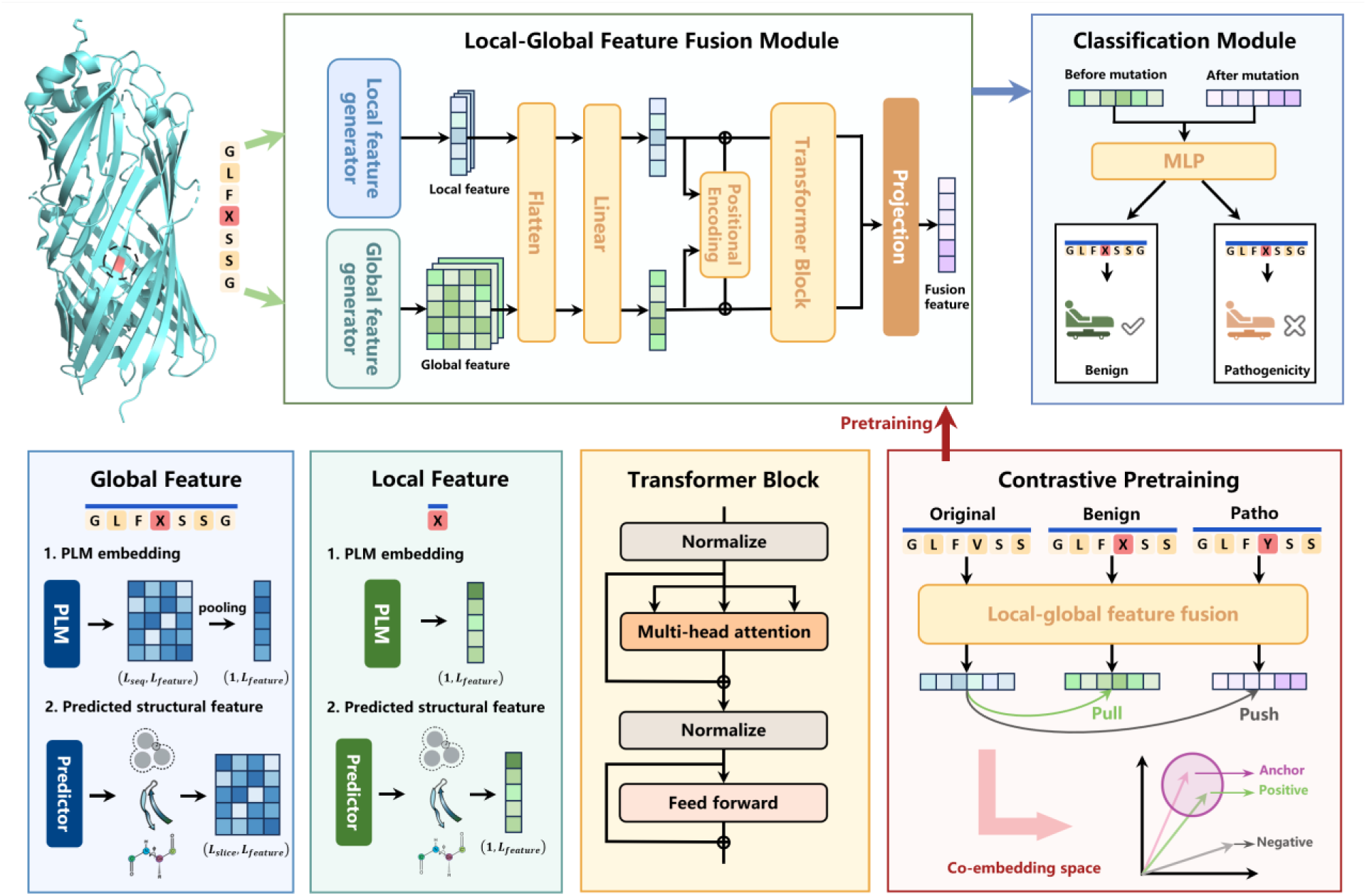
Schematic overview of the Memo-Patho architecture

A distinguishing characteristic of Memo-Patho is its contrastive pretraining strategy, employed prior to fine-tuning. This unsupervised or self-supervised phase enhances the discriminative capability of the feature fusion module. Using triplets comprising an anchor (original sequence), a positive (known benign variant), and a negative (known pathogenic variant), the model is trained via a contrastive loss function. This objective encourages the fused representations of anchor and positive samples to converge in the embedding space, while simultaneously pushing the representations of anchor and negative samples apart. This pretraining step effectively guides the model to discern subtle, pathogenicity-related feature differences within the same protein sequence.

Finally, a multilayer perceptron classifier processes this unified representation to output a final binary classification, predicting whether the mutation is “Benign” or “Pathogenic”. By integrating local and global contexts through a sophisticated fusion mechanism enhanced by contrastive pretraining, Memo-Patho offers a robust approach for accurate, alignment-free pathogenicity prediction in TMPs.

### Efficient mutation pathogenicity prediction using Memo-Patho and comparison with external tools

We developed and evaluated Memo-Patho using two distinct dataset partitioning strategies, termed Mix and Ind datasets, with comprehensive details on data splitting and the two-stage model training (contrastive pre-training followed by binary supervised classification) available in the Methods section. Hyperparameters were tuned using a 10-fold cross-validation approach (cross-validation results are presented in **Supplementary Figures 5-8**), and final model performance was rigorously assessed on independent test sets. We benchmarked Memo-Patho against a suite of contemporary pathogenicity prediction tools, including TransEFVP [35], MutPred2 [24], PROVEAN [38], AlphaMissense [26], PredMutHTP (specific to TMPs) [30], ESNPs&GO [25], and PON-P3 [39]. The methodologies for calculating evaluation metrics and the specific procedures for testing these comparable tools are detailed in the **Supplementary Note** and **Methods**, respectively. Notably, for AlphaMissense and PON-P3, which can output an ‘ambiguous’ classification for predictions near a probability of 0.5, we resolved these into benign or pathogenic categories using a 0.5 probability threshold for consistent comparison, unless explicitly stated otherwise. On the Mix dataset’s independent test set, Memo-Patho achieved a superior accuracy of 0.9317 and a Matthews Correlation Coefficient (MCC) of 0.8618 (**Figure 2a-b, Supplementary Table 1**). Similarly, when evaluated on the Ind dataset’s independent test set, which assesses generalization to proteins unseen during training, Memo-Patho demonstrated robust performance with an accuracy of 0.9146 and an MCC of 0.8209, again outperforming the other tools (**Figure 2a-b, Supplementary Table 2**).

**Figure 2:**
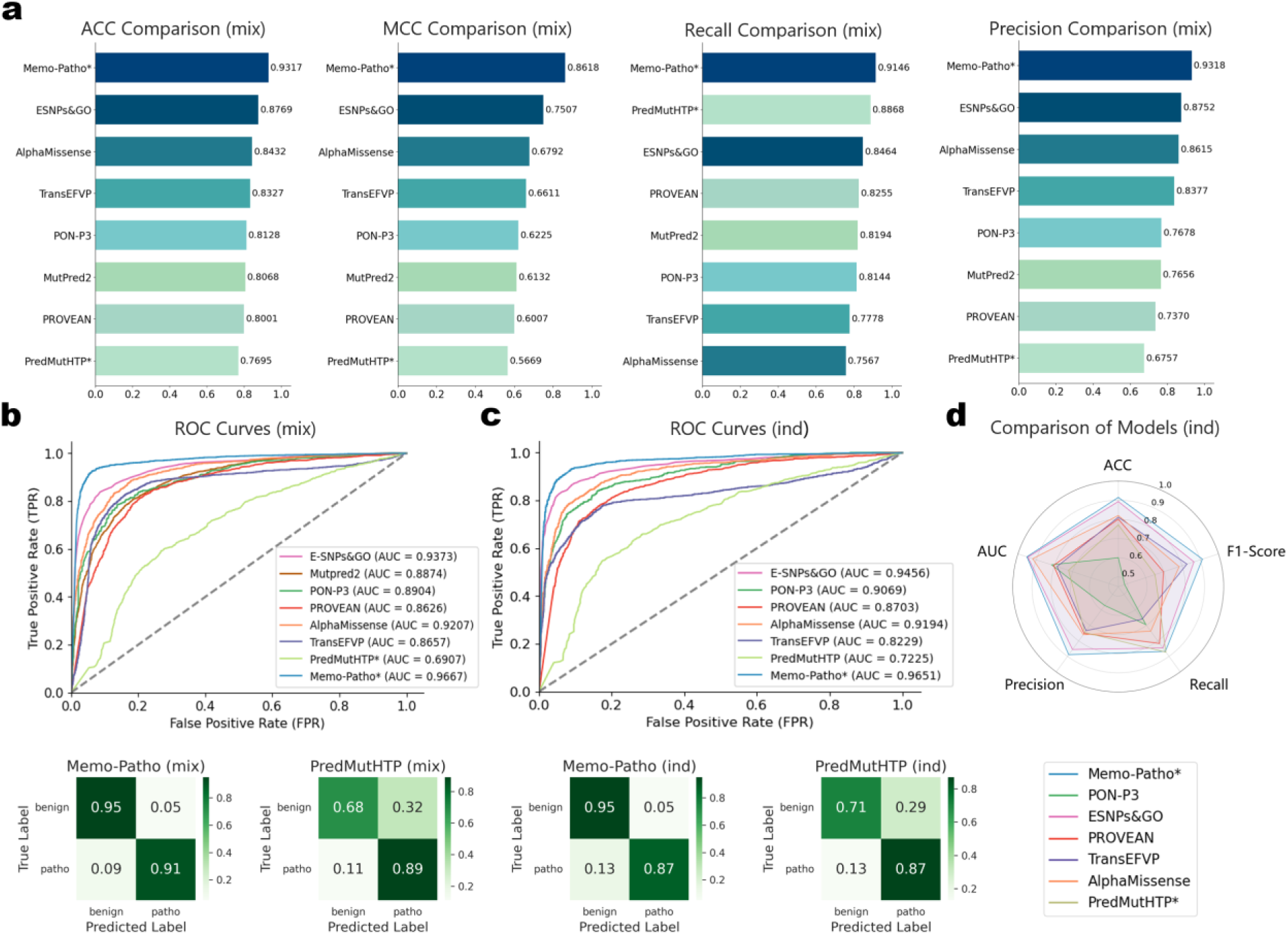
Memo-Patho performance evaluation. **(a-b)** Performance comparison of Memo-Patho with comparable tools on the mix dataset. **(c-d)** Performance comparison of Memo-Patho with comparable tools on the ind dataset. Note that ‘*’ indicates the model is specific to TMPs.

It is important to consider that since we could not ascertain whether our test set entries were excluded from the training sets of these external tools, their reported accuracies might be higher than what would be observed on entirely novel data, a scenario that could further underscore Memo-Patho’s effectiveness. Beyond its predictive accuracy, Memo-Patho offers significant computational efficiency due to its alignment-free architecture. The entire workflow, from input sequence to final classification, can process approximately 5,000 sequences in under 30 minutes on our specified hardware configuration. In contrast, the common preprocessing step of generating multiple sequence alignments using a multi-threaded HHblits tool [40] on the same hardware typically requires around 40 seconds per sequence. This highlights that Memo-Patho’s end-to-end prediction is over a hundred times faster per sequence than this single alignment step frequently required by other methods.

### Enhancing prediction with effective representation learned by contrastive pre-training

To further illustrate the effectiveness of our contrastive pre-training strategy, we first investigated its capacity to learn more discriminative feature representations, beyond its impact on final classification performance. We hypothesized that this pre-training phase would refine the embedding space, making benign and pathogenic variants more separable (specific methodologies for this analysis are detailed in Methods).

To qualitatively assess this, we employed t-SNE [41] for dimensionality reduction and visualization of the feature embeddings. **Figures 3a** and **3c** showcase the feature distributions on the Mix and Ind datasets, respectively, following the application of our contrastive pre-training strategy. These visualizations demonstrate that the contrastive pre-training indeed cultivates a more structured and effective embedding space where samples with different functional impacts begin to form more distinct clusters. Complementing this visual inspection, we conducted a quantitative analysis by evaluating intra-cluster purity. We applied a clustering algorithm to the feature embeddings both before and after contrastive pre-training and then assessed the purity of the resulting clusters with respect to the ground-truth pathogenicity labels. As depicted in **Figures 3b** (for the Mix dataset) and **3d** (for the Ind dataset), there is a marked improvement in intra-cluster purity in the representations obtained after contrastive pre-training compared to the initial, pre-contrastive features. This significant increase in purity provides quantitative evidence that the contrastive pre-training stage successfully enhances the discriminative power of the learned features, aligning them more closely with the inherent biological distinctions between pathogenic and benign mutations.

**Figure 3:**
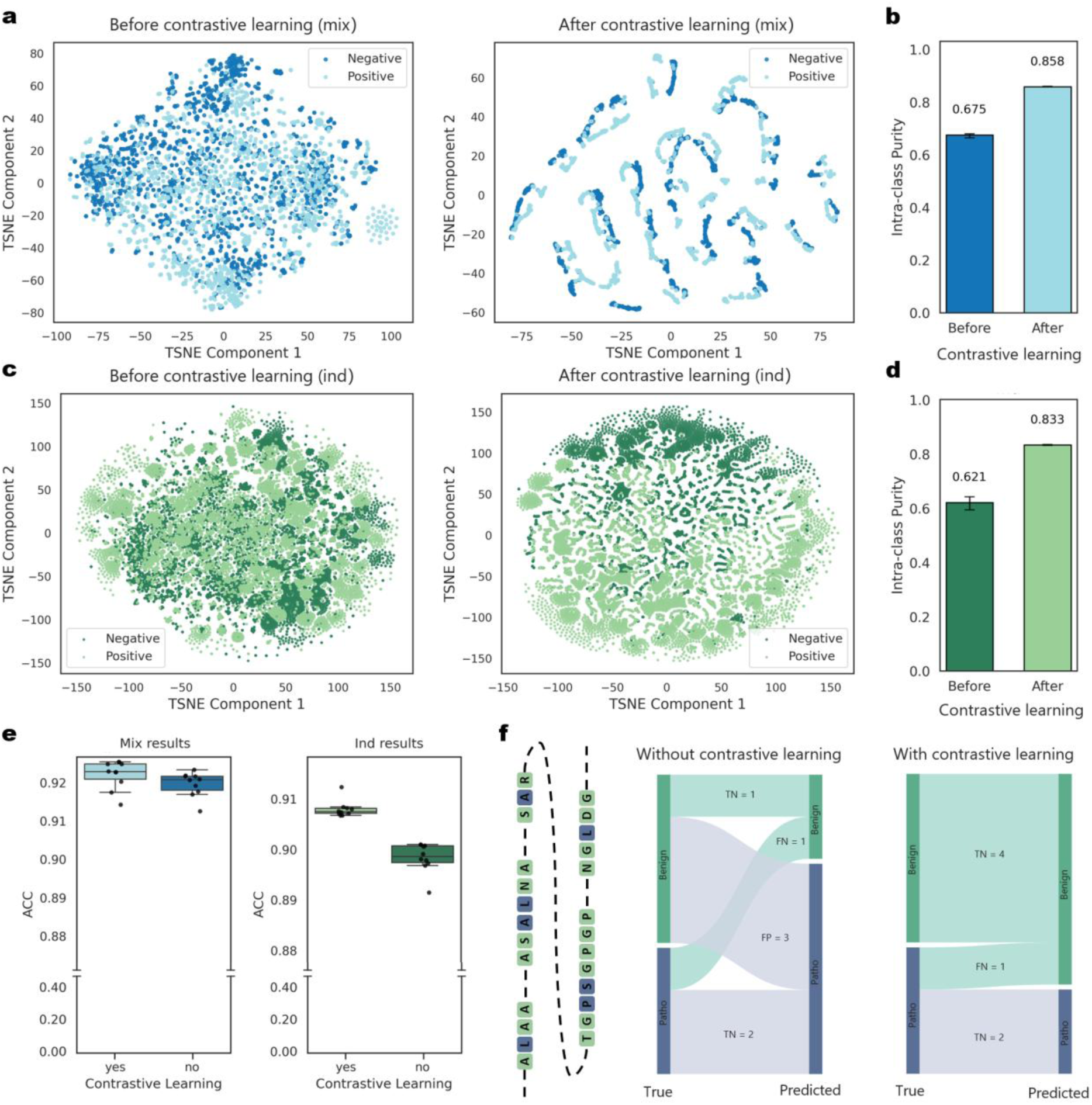
Effectiveness analysis of contrastive learning. **(a)** t-SNE visualization of input and output embeddings on the Mix dataset. **(b)** Comparison of intra-class purity on the Mix dataset before and after contrastive learning. **(c)** t-SNE visualization of input and output embeddings on the Ind dataset. **(d)** Comparison of intra-class purity on the Ind dataset before and after contrastive learning. **(e)** Comparison of Memo-Patho with and without contrastive learning pre-training. **(f)** After contrastive learning, Memo-Patho can effectively handle the complex scenario of varying mutational effects within the same sequence.

Following the analysis of feature representation quality, we directly investigated how contrastive pre-training contributed to the final performance of Memo-Patho. As shown in **Figure 3e**, the inclusion of the contrastive pre-training stage resulted in improved performance metrics during 10-fold cross-validation on both the Mix and Ind datasets. Notably, the performance uplift appeared more significant on the Ind dataset. This more pronounced improvement on the Ind dataset, which is designed to test generalization to entirely unseen protein sequences, likely stems from the pre-training phase encouraging the model to learn more robust and generalizable sequence-level features that capture the subtle differential impacts of mutations on the same protein backbone, rather than potentially overfitting to characteristics of proteins seen during training. This observation suggests that our contrastive pre-training strategy effectively enhances the model’s ability to generalize to novel, previously unencountered sequences.

To further illustrate this practical benefit, **Figure 3f** provides a case study from the Ind dataset’s independent test set, focusing on the protein Q6ZMI3. This protein has three mutations annotated as pathogenic and four as benign in our test data. When Memo-Patho was trained without the contrastive pre-training stage, it incorrectly predicted the outcome for four of these seven variants. However, upon incorporating contrastive pre-training, the model’s accuracy for this specific protein improved dramatically, with only one of the seven variants being misclassified. This example concretely demonstrates the enhanced predictive accuracy and reliability conferred by the contrastive pre-training step.

### Analysis of local-global feature effectiveness revealing the importance of multi-scale features

In this section, we dissected the contributions of local and global features to the overall performance of Memo-Patho. We first conducted ablation studies by comparing the full Memo-Patho model against versions trained using only local features or only global features. **Figures 4a** and **4c** illustrate the 10-fold cross-validation performance of these models on the Mix and Ind datasets, respectively. The results clearly demonstrate that both local features (derived from the immediate vicinity of the mutation) and global features (representing the broader protein context) are individually effective in predicting mutation pathogenicity. Crucially, the combined use of both local and global features in the full Memo-Patho model consistently achieved superior performance, indicating that these two types of features capture complementary information essential for accurate predictions.

**Figure 4:**
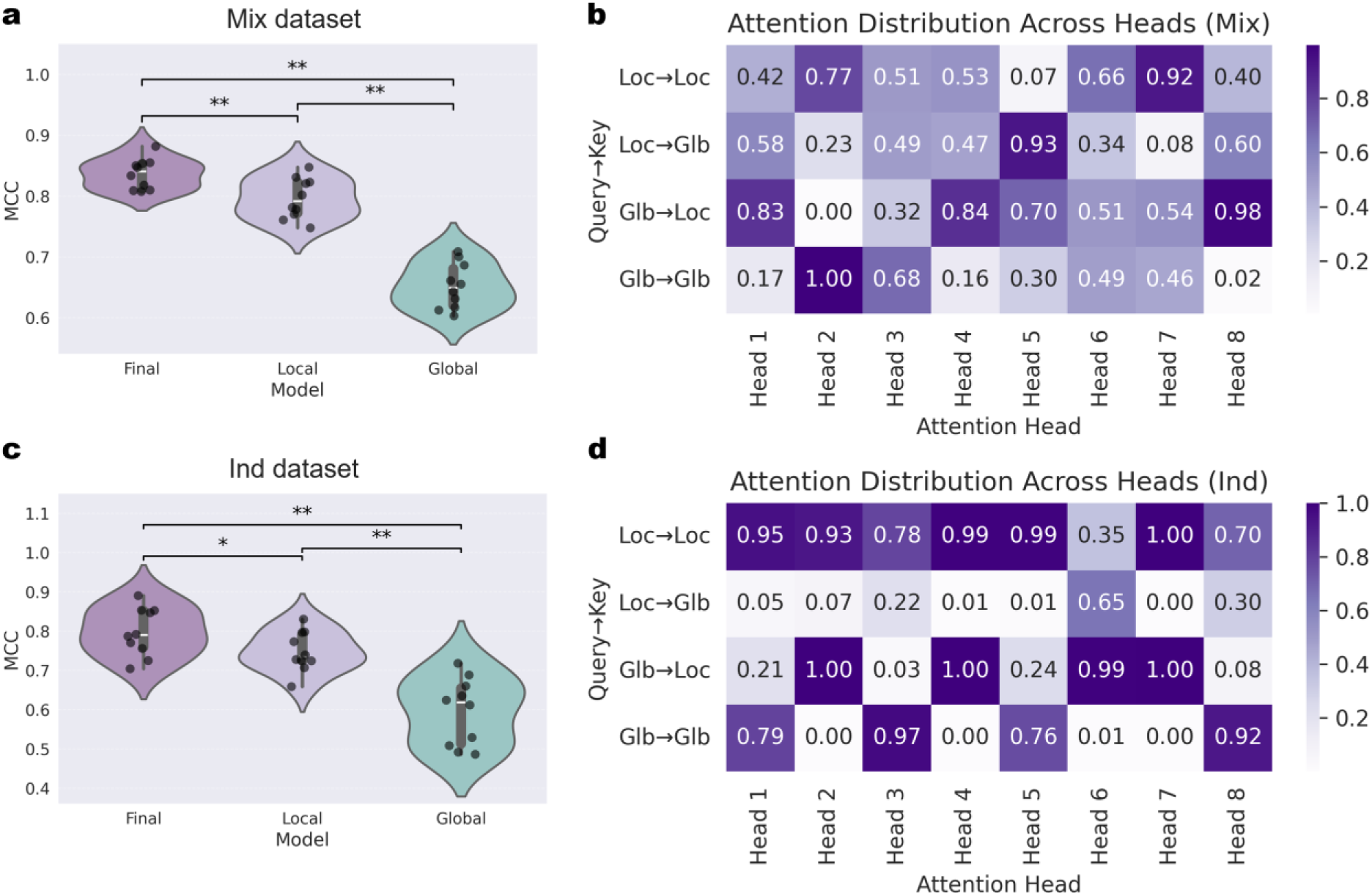
Effectiveness analysis of local-global features. **(a)** Comparison of Memo-Patho with its variants using only local features and only global features on the Mix dataset. **(b)** Attention visualization on the Mix dataset. **(c)** Comparison of Memo-Patho with its variants using only local features and only global features on the Ind dataset. **(d)** Attention visualization on the Ind dataset.

To further understand how local and global features are integrated, we analyzed the attention mechanisms within the feature fusion stage of the mutation processing branch. We extracted the attention coefficients from eight attention heads, examining interactions categorized as local-to-local, local-to-global, global-to-local, and global-to-global. The results, depicted in **Figures 4b** (Mix dataset) and **4d** (Ind dataset), reveal that local features generally command greater attention during the fusion process. This finding aligns with our ablation experiments, which highlighted the strong predictive capacity of local features. Interestingly, we also observed a distinct pattern between the datasets: the Memo-Patho model trained on the Ind dataset (for unseen proteins) placed a comparatively stronger emphasis on local features than the model trained on the Mix dataset. This suggests that when faced with entirely novel sequences, the model prioritizes information directly from the mutation site itself, whereas for sequences it might have encountered previously (as in the Mix dataset, albeit with different mutations), it tends to leverage more of the contextual information from the broader protein environment.

### Enable high-throughput interpretation of protein mutation pathogenicity using Memo-Patho

The alignment-free architecture of Memo-Patho underpins its capacity for rapid processing of extensive sequence datasets, positioning it as an exceptionally suitable tool for large-scale mutational pathogenicity screening. To exemplify this high-throughput capability and further validate our model’s ability to discern biologically relevant patterns, we conducted an exhaustive in silico mutational scan on two proteins selected from our test set: A0A6Q8PFI8, a human protein of 359 amino acids, and B7WPR2, a 451 amino acid human protein known as Transmembrane Serine Protease 3, which plays a role in the inner ear and is associated with deafness. We then investigated the relationship between Memo-Patho’s predictions and sequence conservation. Sequence conservation, reflecting the evolutionary pressure to maintain specific amino acids at positions critical for protein structure or function, serves as an independent biological indicator of residue importance. Mutations at highly conserved sites are generally more likely to be deleterious [42]. The detailed methodology for calculating conservation scores and performing this comparison is provided in Methods. For protein A0A6Q8PFI8, the distribution of site-specific conservation scores (**Figure 5a**) and Memo-Patho’s average predicted pathogenicity scores per residue (**Figure 5b**) demonstrate a compelling visual correspondence. This relationship was quantified with a Pearson correlation coefficient of 0.69 (**Figure 5c**), signifying a strong positive correlation. Similar strong correlations and visual concordance were observed for B7WPR2 (**Figures 5d-f**). The consistent and strong positive correlation between Memo-Patho’s predictions and this fundamental measure of residue importance (conservation) strongly suggests that our model captures genuine biological signals related to the functional or structural indispensability of residues. This finding enhances confidence in Memo-Patho’s ability to identify potentially critical mutations, even in the absence of extensive experimental data for every possible variant.

**Figure 5:**
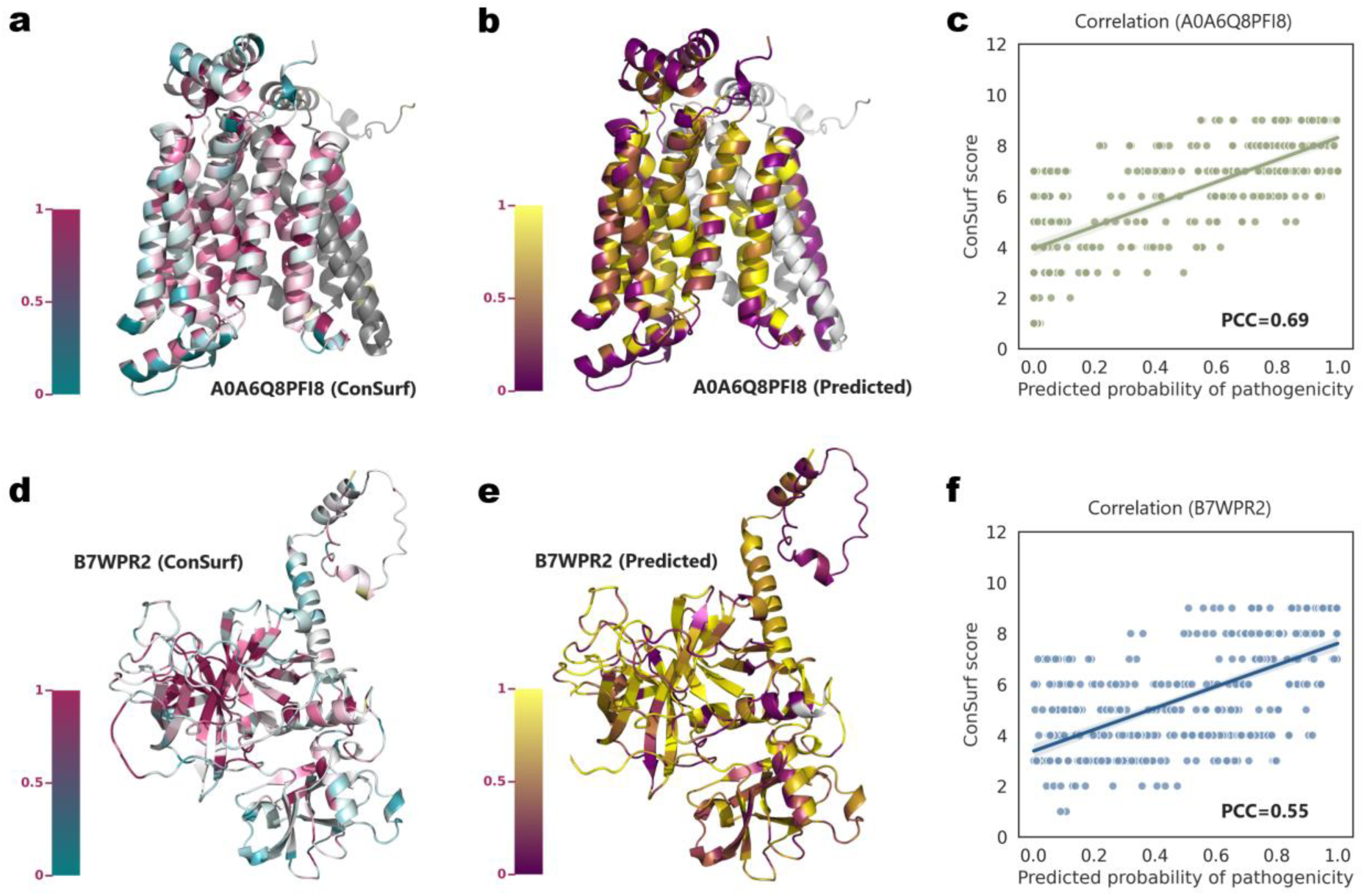
Correlation analysis with conservation. **(a-b)** Visualization of conservation scores and Memo-Patho prediction scores on A0A6Q8PFI8. **(c)** Correlation plot of conservation scores versus Memo-Patho prediction scores on A0A6Q8PFI8. **(d-e)** Visualization of conservation scores and Memo-Patho prediction scores on B7WPR2. **(f)** Correlation plot of conservation scores versus Memo-Patho prediction scores on B7WPR2.

The comprehensive mutational scan for A0A6Q8PFI8 and B7WPR2 generated a dataset of 15,390 unique amino acid substitutions. Memo-Patho processed this entire dataset, delivering pathogenicity predictions for all variants in under 80 minutes on our standard hardware. Such rapid, exhaustive screening, particularly when its outputs align well with biological indicators like sequence conservation, underscores Memo-Patho’s significant practical utility. It empowers researchers to systematically probe entire protein mutational landscapes with high throughput, thereby accelerating the identification of critical regions, the prioritization of variants for experimental validation, and the exploration of previously uncharacterized segments of the mutational space for novel insights into disease mechanisms or protein function.

### Memo-Patho unveils functional impact of critical KCNQ1 variants

To further validate Memo-Patho’s effectiveness and its ability to generalize to entirely independent data, we conducted an additional round of testing on a novel dataset of variants in the KCNQ1 protein. This dataset was recently established by Brewer et al. (published February 19, 2025) and comprises 93 variants [43] sourced from databases such as HGMD [44], including those with known, controversial, or previously undetermined clinical significance. The KCNQ1 gene encodes a critical voltage-gated potassium channel, and its loss-of-function is the predominant cause of Type 1 Long QT Syndrome (LQTS1), one of the most prevalent inherited cardiac arrhythmogenic disorders [45]. The study by Brewer et al. involved a comprehensive suite of experimental analyses, meticulously probing the functional, trafficking, and biophysical properties of these variants to definitively ascertain their pathogenicity. From this dataset, we excluded variants that overlapped with our own training and initial test sets, resulting in a stringent novel test set of 55 KCNQ1 mutation records. The structural distribution of these 55 mutations on the KCNQ1 protein is depicted in **Figure 6a**, while their distribution across various structural regions and the composition of their assigned pathogenic/benign labels are detailed in **Figures 6b** and **6c**, respectively.

**Figure 6:**
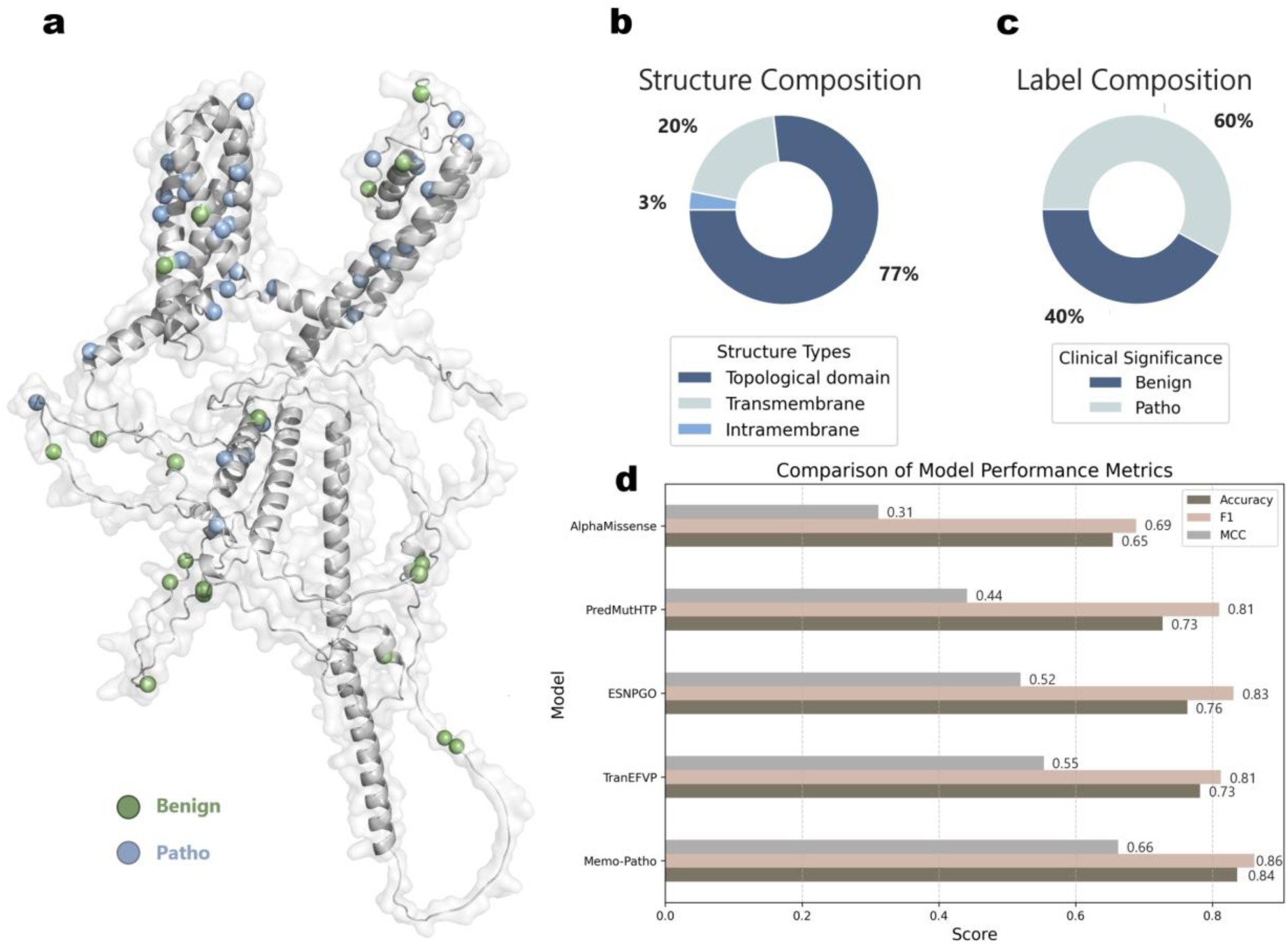
Performance comparison on the novel dataset. **(a)** Visualization of mutations on the KCNQ1 and novel datasets (green indicates benign mutations, blue indicates pathogenic mutations). **(b)** Distribution of topological structure types for KCNQ1. **(c)** Label distribution on the novel mutation dataset. **(d)** Predictive performance comparison of Memo-Patho with other state-of-the-art tools.

When evaluated on this challenging KCNQ1 dataset, Memo-Patho demonstrated superior predictive performance compared to other contemporary tools, achieving an accuracy of 0.84 and an MCC of 0.66 (**Figure 6d**). This strong performance on a rigorously characterized, clinically highly relevant, and entirely novel protein dataset effectively underscores Memo-Patho’s robust generalization capabilities. Such results indicate that our model is not merely fitting to its training data but has learned underlying principles of variant pathogenicity, enabling its reliable application in diverse and demanding clinical and research scenarios. The ability to accurately classify variants in a well-studied disease gene like KCNQ1, whose variants were experimentally validated with such depth by Brewer et al., is particularly significant. It suggests Memo-Patho can serve as a powerful adjunctive tool for prioritizing variants of uncertain significance (VUS) for further experimental investigation, potentially accelerating diagnostic workflows and improving the understanding of disease mechanisms in clinically actionable genes.

## Discussion

Accurately predicting protein mutation pathogenicity is pivotal for genomics and personalized medicine. Existing methods often struggle with generalization, reliance on time-consuming MSAs, or rapid large-scale screening. Our study introduces Memo-Patho, an alignment-free deep learning framework integrating diverse sequence and predicted structural features via a novel two-stage training process, including contrastive pre-training, designed for accurate, generalizable, and computationally efficient predictions.

Memo-Patho directly addresses key challenges in variant interpretation. Its alignment-free nature circumvents the MSA bottleneck, while innovative contrastive pre-training sensitizes the model to subtle differential mutational effects on identical protein backbones. Combined with rich features from potent protein language models (ESM2, ProtT5) and predicted structural properties (NetSurfP-3.0), Memo-Patho learns robust, generalizable representations, making it well-suited for large-scale genomic screening and exploring uncharacterized mutational landscapes.

The development of robust predictors like Memo-Patho holds particular significance for TMPs. These proteins are integral to cellular physiology, serving as crucial drug targets and frequently implicated in disease. However, pathogenic annotations for TMPs are relatively scarce compared to globular proteins, potentially leading to their underrepresentation in general predictors’ training data. Consequently, such models may not fully capture unique TMP constraints, leading to an information dilution where TMP-specific signals are overshadowed. Memo-Patho’s strong performance, achieved without MSA reliance and bolstered by effective feature learning, offers a valuable approach to address this gap, providing more focused insights into variation within this critical protein superfamily.

Memo-Patho’s strategies yield state-of-the-art performance and robust generalization on our TMP-centric internal Mix and Ind datasets. This capacity was further affirmed by its superior performance on a challenging, externally curated KCNQ1 (a key TMP ion channel) variant dataset. Given KCNQ1’s pivotal role in cardiac function, Memo-Patho’s success on this dataset underscores its potential for real-world clinical applicability, especially for TMP-related disorders.

Our analyses confirmed the synergistic value of local and global features. Attention mechanisms generally prioritized local features, a focus that intensified for novel proteins (’Ind’ dataset), suggesting a sophisticated, context-aware integration strategy. This aligns with the refined feature space achieved through contrastive pre-training, enhancing the model’s ability to distinguish mutational effects.

Memo-Patho’s biological relevance is further supported by a strong positive correlation (e.g., Pearson coefficient of 0.69 for A0A6Q8PFI8, with similar findings for TMPRSS3/B7WPR2, as detailed in Figures 5a-f) between its predictions and evolutionary conservation scores. This alignment suggests it captures intrinsic features of deleterious impacts, vital for TMPs and less-studied proteins.

A key advantage is Memo-Patho’s exceptional computational efficiency due to its alignment-free design, processing thousands of sequences orders of magnitude faster than MSA-based approaches. This renders it highly practical for comprehensive in silico mutational scans, including those focused on the transmembrane proteome.

Despite its robust performance and notable advantages, Memo-Patho shares limitations common to many complex deep learning models. A primary area for future development is enhancing interpretability. While attention mechanisms offer some high-level insights into feature importance, the precise mechanistic basis for individual predictions can be opaque, making it challenging to fully deconstruct the model’s decision-making process. Future work could explore the integration of causal inference methodologies to more rigorously identify determinant features and analyze model predictions in conjunction with diverse, known molecular mechanisms of pathogenicity, thereby aiming to build a more transparent understanding of how different mutational patterns are recognized. Secondly, Memo-Patho’s knowledge is derived from experimentally validated mutations in public databases like ClinVar. The methodical, resource-intensive nature of variant characterization means current datasets may not uniformly cover all mutations, contexts, or pathogenic pathways, potentially introducing latent biases. This could affect performance on rare or novel variants underrepresented in training data. We anticipate that advancements in high-throughput experimental technologies will be crucial for generating more comprehensive datasets, thereby mitigating biases and refining models like Memo-Patho alongside our expanding experimental understanding.

Overall, Memo-Patho represents a significant advancement in predicting mutation pathogenicity. By thoughtfully integrating advanced deep learning techniques—including a novel contrastive pre-training scheme and attention mechanisms—with diverse biological features in an alignment-free framework, it offers a powerful, generalizable, and exceptionally efficient tool. Its particular strengths in handling TMPs address a notable need in the field. As models like Memo-Patho evolve and are prospectively validated, they are poised to become indispensable in genomic research, clinical variant interpretation, and advancing precision medicine.

## Methods

### Data preparation and preprocessing

Human transmembrane protein sequences were obtained from the UniProt Knowledgebase (UniProtKB, accessed January 20, 2025), a central hub for curated protein sequence and functional information [46]. Variant annotations were sourced from ClinVar, a public archive documenting relationships between human genetic variations and observed health statuses [47]. For our analysis, ClinVar’s ‘Likely benign’ and ‘Benign’ labels were consolidated into a ‘benign’ category, while ‘Likely pathogenic’ and ‘Pathogenic’ labels were grouped as ‘pathogenic’. Mapping these annotations to the protein sequences initially yielded 13,226 proteins associated with 38,822 pathogenic and 48,347 benign variants. We then performed a quality control step, removing annotations where the reported wild-type residue in ClinVar conflicted with the residue in the corresponding UniProt reference sequence. This filtering process resulted in a final dataset of 81,884 curated mutation annotations used for subsequent analyses.

### Extraction of global and local Features

To capture comprehensive sequence information, we utilized embeddings derived from two state-of-the-art PLMs: ESM2 and ProtT5. ESM2 is a deep transformer-based model developed by Meta AI, pre-trained on vast quantities of protein sequences to learn underlying biological properties [48]. ProtT5 is an encoder-decoder transformer model from the Rost Lab, also trained on large protein datasets and adept at capturing sequence-level patterns relevant to structure and function [31]. Leveraging both models is advantageous, as prior studies suggest they can offer complementary insights into protein characteristics [35, 49]. Specific details of the embedding generation are available in the Supplementary Note. For features derived from ESM2, which produces embeddings with dimensions (sequence length × 2560), the local feature was defined as the specific embedding vector corresponding to the mutated residue position (1 × 2560). The global feature was obtained by applying mean pooling across the entire sequence’s embedding vectors, resulting in a fixed-size representation (1 × 2560). An analogous procedure was applied to the ProtT5 embeddings (output dimensions: sequence length × 1024). The local feature was extracted directly from the mutation site (1 × 1024), and the global feature was generated via mean pooling over the full sequence embedding (1 × 1024). The formula is as follows:

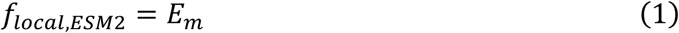

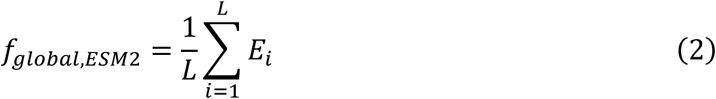

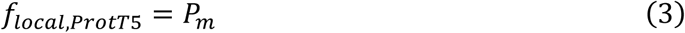

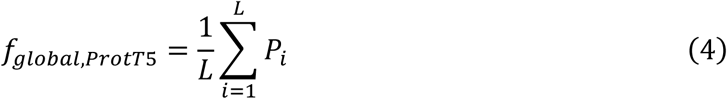

where *S* is the protein sequence, *L* is its length, *m* is the position index of the mutated residue (1 ≤ *m* ≤ *L*), *ESM*2(*S*) is the *L* × 2560 embedding matrix output by the ESM2 model for sequence *S*, *ProtT*5(*S*) is the *L* × 1024 embedding matrix output by the ProtT5 model for sequence *S*, *E*_*i*_ ∈ *R*^1×2560^ is the row vector within *ESM*2(*S*) corresponding to the *i*-th residue, and *P*_*i*_ ∈ *R*^1×1024^ is the row vector within *ProtT*5(*S*) corresponding to the *i*-th residue.

Furthermore, we incorporated predicted structural features into our analysis, including relative solvent accessibility (RSA), absolute solvent accessibility (ASA), secondary structure (SS), backbone dihedral angles, and intrinsic disorder. These features were included as numerous studies have indicated direct or indirect associations between local structural characteristics and variant pathogenicity [50, 51]. To obtain these predictions, we utilized NetSurfP-3.0 [52], a prediction tool that leverages PLM embeddings, thereby avoiding computationally expensive multiple sequence alignments, and has been specifically fine-tuned on structural annotation tasks. NetSurfP-3.0 outputs a consolidated 16-dimensional feature vector per residue summarizing these predicted structural properties. For each variant, the local structural feature was defined as this 16-dimensional vector corresponding precisely to the mutated residue position. To represent the structural context, a global feature was constructed by considering a window of residues spanning four positions upstream and four positions downstream of the mutation site (9 residues in total). The 16-dimensional feature vectors for these 9 residues were then aggregated to form a 9×16 matrix representing the local structural environment. In cases where the mutation occurred near the sequence termini and the full 9-residue window could not be formed, the missing positions in the matrix were padded with a value of -1. The formula is as follows:

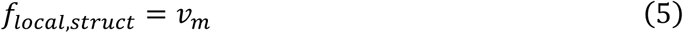

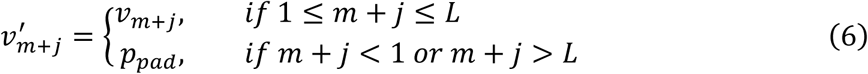

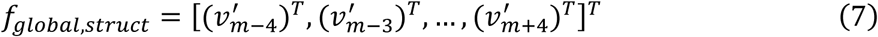

where *NSP*(*S*) is the *L* × 16 output matrix from NetSurfP-3.0 for sequence S, *v*_*i*_ ∈ *R*^1×16^ is the *i*-th row vector of *NSP*(*S*) representing the 16-dimensional structural feature vector for the *i* -th residue, and *p*_*pad*_ ∈ *R*^1×16^ is a padding vector consisting entirely of -1 values.

### Dataset construction and partitioning

To comprehensively demonstrate the effectiveness and generalization capability of our strategy, we constructed two distinct datasets and trained two corresponding versions of our model, Memo-Patho. The first dataset, termed the Mix dataset, was created by partitioning the data at the level of individual mutation entries. We randomly sampled 10% of all mutation records to form the independent test set, using the remaining data for the training and validation sets. Consequently, protein sequences appearing in the Mix test set might also be present in the training/validation data, albeit associated with different, unseen mutations. The model trained on this dataset serves as our final released version and consists of 47,677 training/validation entries and 5,298 test entries. The second dataset, termed the ‘Ind’ dataset, was partitioned at the protein sequence level. Here, 10% of the unique protein sequences, along with all their associated mutations, were randomly allocated to the independent test set. This ensures that no protein sequence in the Ind test set was encountered during model training or validation, providing a more stringent evaluation of the model’s ability to generalize to novel proteins. The Ind dataset comprises 48,135 training/validation entries and 4,840 test entries. The specific distributions across training, validation, and test sets for both the Mix and Ind datasets are detailed in **Supplementary Figure 1-4**.

### Contrastive pre-training

We incorporated a contrastive pre-training stage specifically designed to train the model to distinguish between representations associated with benign versus pathogenic outcomes originating from the same wild-type protein sequence. To achieve this, we first constructed a dataset of contrastive triplets using our training data. For each protein sequence present in the training set with both benign and pathogenic annotations, we defined: (i) the anchor (A) as the representation derived from the original (wild-type) sequence features; (ii) the positive sample (P) as the representation corresponding to the sequence harboring a mutation classified as benign; and (iii) the negative sample (N) as the representation corresponding to the sequence harboring a mutation classified as pathogenic. This process resulted in a pre-training dataset comprising 213,031 triplets. During the pre-training phase, the learning objective was to adjust the model parameters to minimize the distance between the fusion features of the anchor and positive samples (A-P), while simultaneously maximizing the distance between the fusion features of the anchor and negative samples (A-N). This encourages the model to learn discriminative representations sensitive to the functional impact of mutations within the context of an identical protein backbone.

### Memo-Patho training methods and settings

In the training phase of Memo-Patho, we defined a sequence receptive field of 501 residues centered on the mutation site. This means that for feature extraction and model input, we considered the mutated residue along with 250 residues upstream and 250 residues downstream. Utilizing such a fixed-length context window is a common practice when applying deep learning to protein sequences [53, 54]. This ensures consistent input dimensionality for the model and balances the capture of relevant local sequence information against the computational complexity and memory demands that can arise from processing entire, potentially very long, protein sequences. To ensure robust evaluation and mitigate potential dataset biases, we employed a 10-fold cross-validation strategy during training. In this approach, the relevant training/validation dataset was randomly partitioned into 10 equally sized folds. The model was then trained 10 independent times. In each iteration, one distinct fold served as the validation set, while the remaining 9 folds were used for training. For model optimization, we utilized the Cross-Entropy Loss function [55] with the Adam optimizer [56]. Key hyperparameters were set as follows: a batch size of 32 and an initial learning rate of 1 × 10^−3^. The Memo-Patho model was implemented using the PyTorch framework [57], and all training and testing experiments were conducted on a workstation equipped with three Nvidia RTX 3090 GPUs and 256GB of system RAM.

### Evaluation workflow for comparable tools

We evaluated its performance against several established pathogenicity prediction tools, following recommended or feasible usage protocols for each. (i) AlphaMissense: Predictions were obtained programmatically by querying the official AlphaMissense web API for specific residue positions (e.g., https://alphamissense.hegelab.org/hotspot?uid=[UniProtID]&resi=[Position]). (ii) TransEFVP: This method was reproduced locally based on its described SVM+PCA approach, applying a fixed input sequence context window of 201 residues (±100 residues around the mutation site). (iii) E-SNPs&GO: Predictions were generated via its web server (https://esnpsandgo.biocomp.unibo.it/), which accepts full sequences without context length restrictions and supports batch input of up to 1000 mutations, including multiple mutations per protein in a single submission. (iv) PredMutHTP: Similar to E-SNPs&GO, this tool was evaluated using its web server (https://www.iitm.ac.in/bioinfo/PredMutHTP/pred.php), accepting full sequences and batch inputs of up to 1000 mutations. (v) PROVEAN: Predictions were obtained using the human species setting on its batch submission web server (http://provean.jcvi.org/protein_batch_submit.php?species=human), which processes full sequences. (vi) MutPred2: Evaluated via its web server (http://mutpred.mutdb.org/#qform), this tool accepts full sequences but limits batch submissions to a maximum of 100 mutations and a total sequence length not exceeding 35,000 residues per job.

### Contrastive pre-training effectiveness analysis

To evaluate the impact of the contrastive pre-training stage on feature representation, we conducted both qualitative visualization and quantitative purity analyses. For dimensionality reduction and subsequent visualization, we applied the t-Distributed Stochastic Neighbor Embedding (t-SNE) algorithm to two types of feature sets: (i) the concatenated input features fed into the model, and (ii) the ‘fusion embeddings’ output by the model after the contrastive pre-training phase. These visualizations were generated using the TSNE implementation from the Python scikit-learn library, adhering to its default parameter settings. For the quantitative intra-cluster purity analysis, we first employed the K-means clustering algorithm, also from scikit-learn, to group the feature embeddings (both before and after contrastive pre-training). To ensure that our purity assessment was robust and not an artifact of a specific number of clusters, we systematically varied the number of clusters (k) for KMeans from 20 to 70, examining the trend of purity improvement across this range.

### Correlation analysis of Memo-Patho prediction scores with conservation

To investigate the effectiveness of Memo-Patho for large-scale mutational space screening and to assess the biological relevance of its predictions, we analyzed the correlation between its site-specific average prediction scores and evolutionary conservation scores. For this analysis, we selected two proteins, A0A6Q8PFI8 and TMPRSS3, and computationally enumerated all possible single amino acid substitutions for each position within these sequences, generating a total of 15,390 unique mutation entries. Site-specific conservation scores were determined using the ConSurf web server (available at https://consurf.tau.ac.il/) [58]. The ConSurf methodology involves an initial search for homologous sequences, followed by the construction of a multiple sequence alignment using MAFFT [59]. Subsequently, an empirical Bayesian algorithm is employed to calculate evolutionary conservation scores based on the phylogenetic relationships between the sequences, classifying each site into one of nine discrete grades, where a score of 1 indicates the most variable (least conserved) positions and a score of 9 signifies the most conserved positions. After obtaining pathogenicity predictions from Memo-Patho for all 15,390 generated mutations, we calculated a single representative prediction score for each residue position by averaging Memo-Patho’s output scores for all possible mutations occurring at that specific site. Visualizations presented in **Figure 5**, such as mapping these scores onto protein structures, were rendered using PyMOL [60]. The Pearson correlation coefficient, quantifying the linear relationship between the site-specific ConSurf conservation scores and Memo-Patho’s average site-specific prediction scores, was computed using the pearsonr function from the scipy.stats module in Python.

### Novel dataset validation

In the evaluation conducted on the novel KCNQ1 dataset (Brewer et al., published February 19, 2025), we employed the identical testing strategies and computational pipelines as those used for our internal datasets (Mix and Ind). This ensured consistency in feature generation, model inference, and the application of evaluation metrics. Annotation information regarding the KCNQ1 protein, including sequence features and regional demarcations (as referenced for **Figure 6b**), was obtained from the UniProt database. For structural visualization purposes, specifically to map the spatial distribution of the 55 tested variants (as shown in **Figure 6a**), we utilized the full-length protein structure model of KCNQ1 as predicted by AlphaFold2 [27]. This choice was made to ensure a complete and uniform structural template for comprehensive visualization of variant locations.

## Supporting information

Supplemental Online Content

## Data availability

Sequence information was obtained from UniProt (https://www.uniprot.org/), and variant annotations were sourced from ClinVar (https://www.ncbi.nlm.nih.gov/clinvar/). Sequence conservation results were derived from ConSurf, with the outputs downloaded from https://consurf.tau.ac.il/. AlphaMissense prediction results were retrieved from https://alphamissense.hegelab.org via its API. KCNQ1 mutation annotation data were directly extracted from the supplementary data of the article available at https://www.pnas.org/doi/full/10.1073/pnas.2412971122. The datasets and features preprocessed in this study are available at https://github.com/RoarBoil/Memo-Patho.

## Code availability

The data and source code of EMO are available at https://github.com/RoarBoil/Memo-Patho.

## Author Contributions

Y.B. and Z.L. conceived the idea of this research and wrote the manuscript. Y.B. processed the data, implemented the predictor, and tuned the model. F.Z. conducted model comparisons and testing, and participated in writing the manuscript. W.L. and J.H. participated in the conservation analysis. G.N.L. supervised the research and reviewed the manuscript. All authors contributed to the article and approved the submitted version.

## Acknowledgments

This work was supported by grants from STI 2030—Major Projects (no. 2022ZD0209100), the Medical-Engineering Cross Foundation of Shanghai Jiao Tong University (nos. YG2022ZD026 and YG2023ZD27), and SJTU Trans-med Awards Research (no. 20220103)

