## Supplemental Online Content for "Memo-Patho: Bridging Local-Global Transmembrane Protein Contexts with Contrastive Pretraining for Alignment-Free Pathogenicity Prediction"

#### Supplementary Figures

1) **Supplementary Figure 1.** Label distribution across datasets.
2) **Supplementary Figure 2.** Mutation position distribution by clinical significance across datasets.
3) **Supplementary Figure 3.** Sequence length distribution across datasets.
4) **Supplementary Figure 4.** Confusion matrix of Memo-Patho's 10-fold cross-validation results on the Mix dataset.
5) **Supplementary Figure 5.** Radar chart of Memo-Patho's 10-fold cross-validation results on the Mix dataset.
6) **Supplementary Figure 6.** Radar chart of Memo-Patho's 10-fold cross-validation results on the Mix dataset.
7) **Supplementary Figure 7.** Confusion matrix of Memo-Patho's 10-fold cross-validation results on the Ind dataset.
8) **Supplementary Figure 8.** Radar Chart of Memo-Patho's 10-fold cross-validation results on the Ind dataset.

#### Supplementary Notes

1) PLMs embedding generation.
2) Calculation methods for evaluation metrics.

#### Supplementary Tables

1) ***Supplementary Table 1.*** *Performance evaluation between baseline models on Mix dataset*
2) ***Supplementary Table 2.*** *Performance evaluation between baseline models on Ind dataset*
3) ***Supplementary Table 3.*** *Performance evaluation between baseline models on novel dataset*

### Supplementary Figure

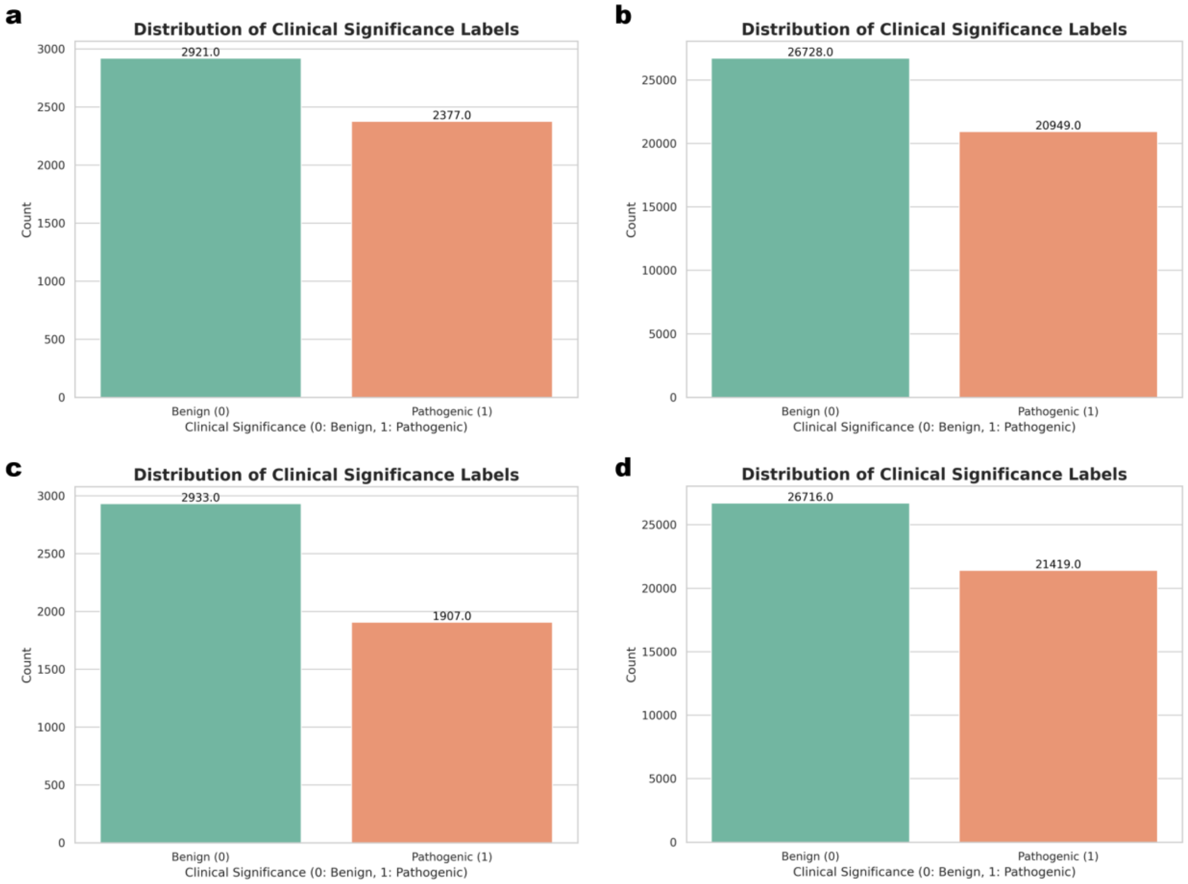


**Supplementary Figure 1.** Label distribution across datasets. **(a)** Mix test set. **(b)** Mix train set. **(c)** Ind test set. **(d)** Ind train set.

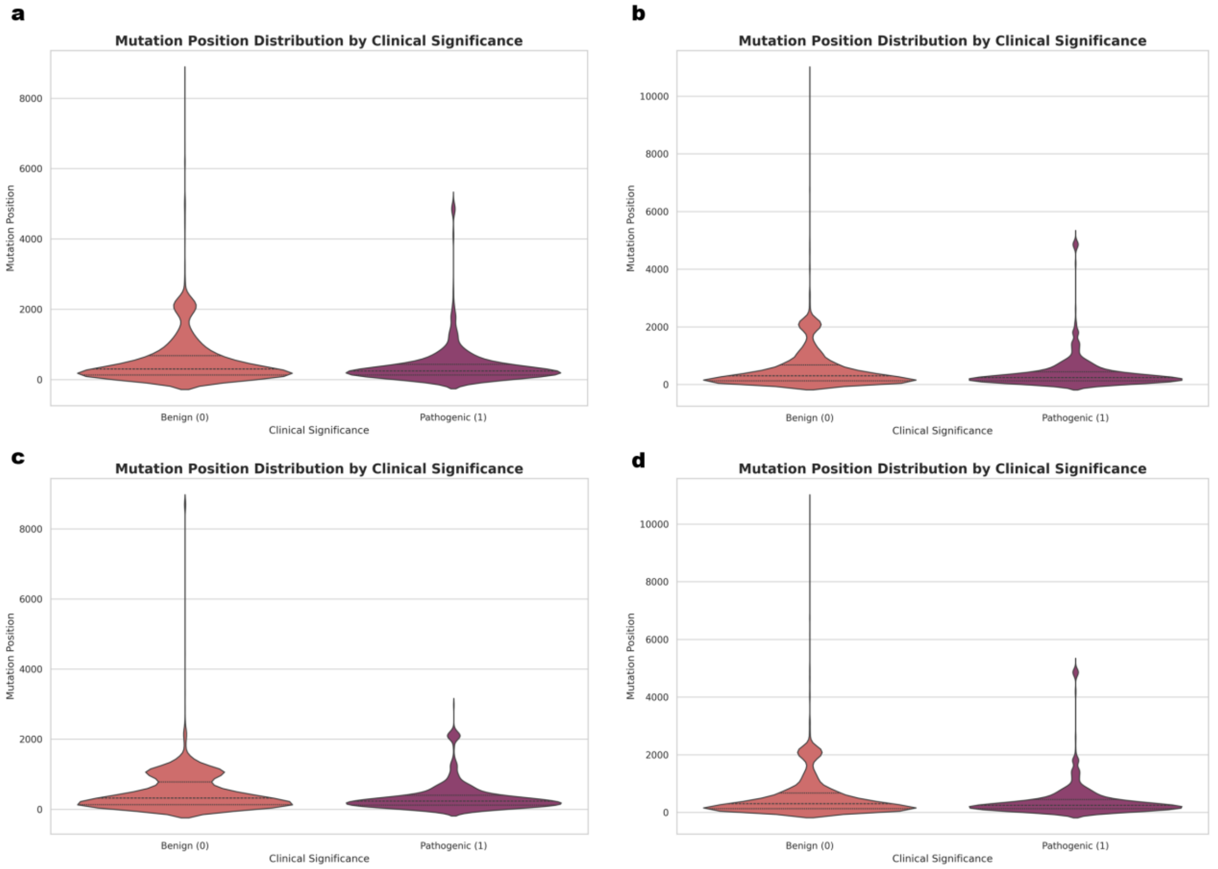


**Supplementary Figure 2.** Mutation position distribution by clinical significance across datasets. **(a)** Mix test set. **(b)** Mix train set. **(c)** Ind test set. **(d)** Ind train set.

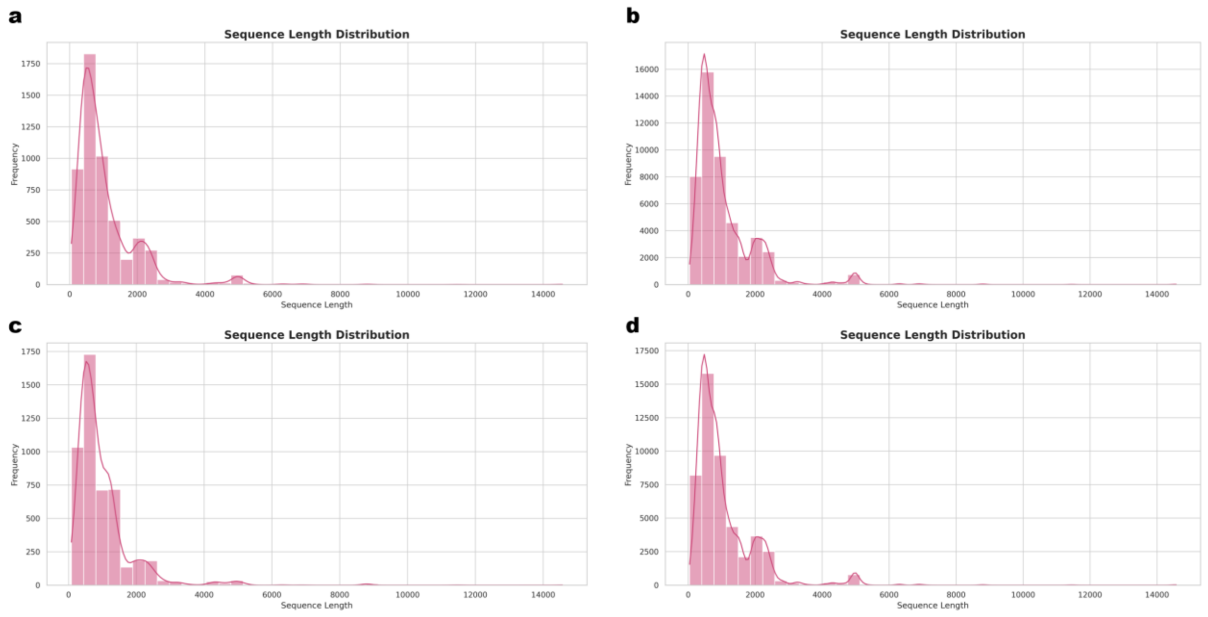


**Supplementary Figure 3.** Sequence length distribution across datasets. **(a)** Mix test set. **(b)** Mix train set. **(c)** Ind test set. **(d)** Ind train set.

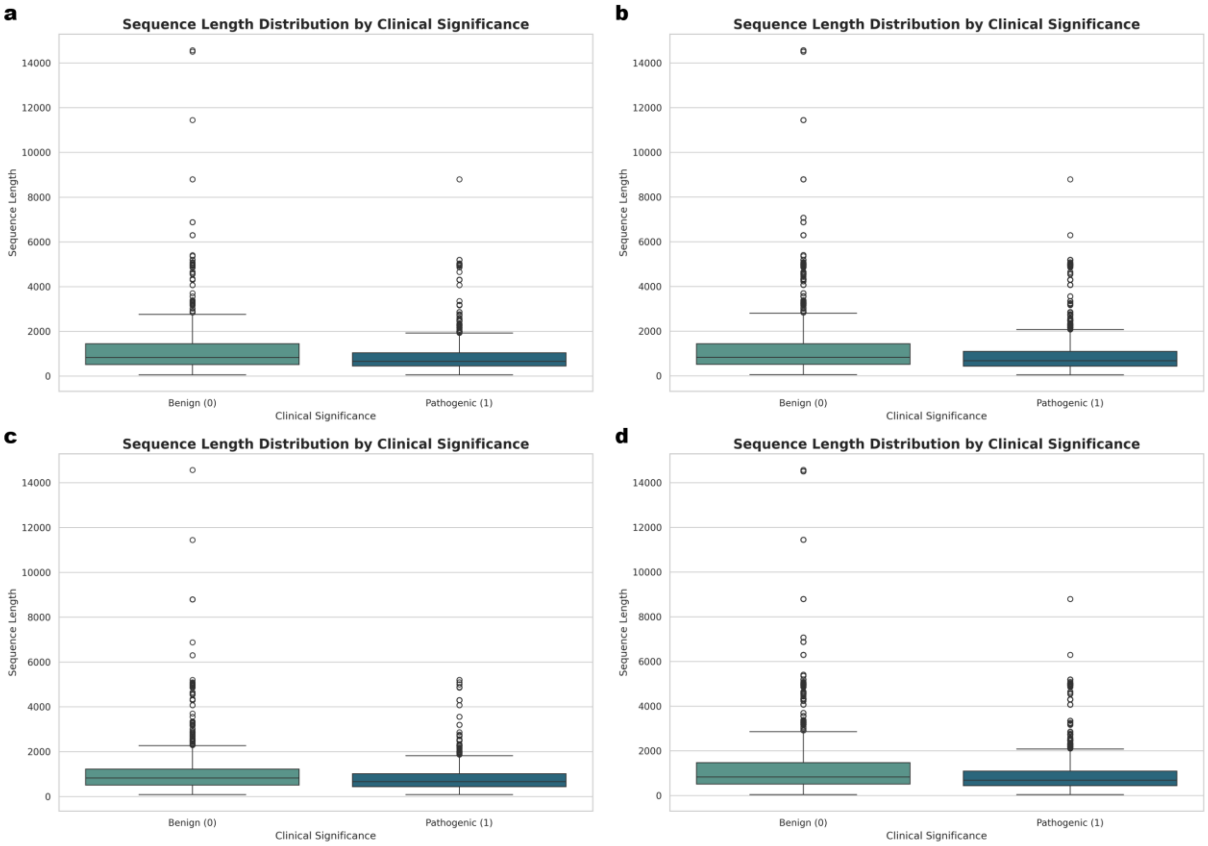


**Supplementary Figure 4.** Sequence length distribution by clinical significance across datasets. **(a)** Mix test set. **(b)** Mix train set. **(c)** Ind test set. **(d)** Ind train set.

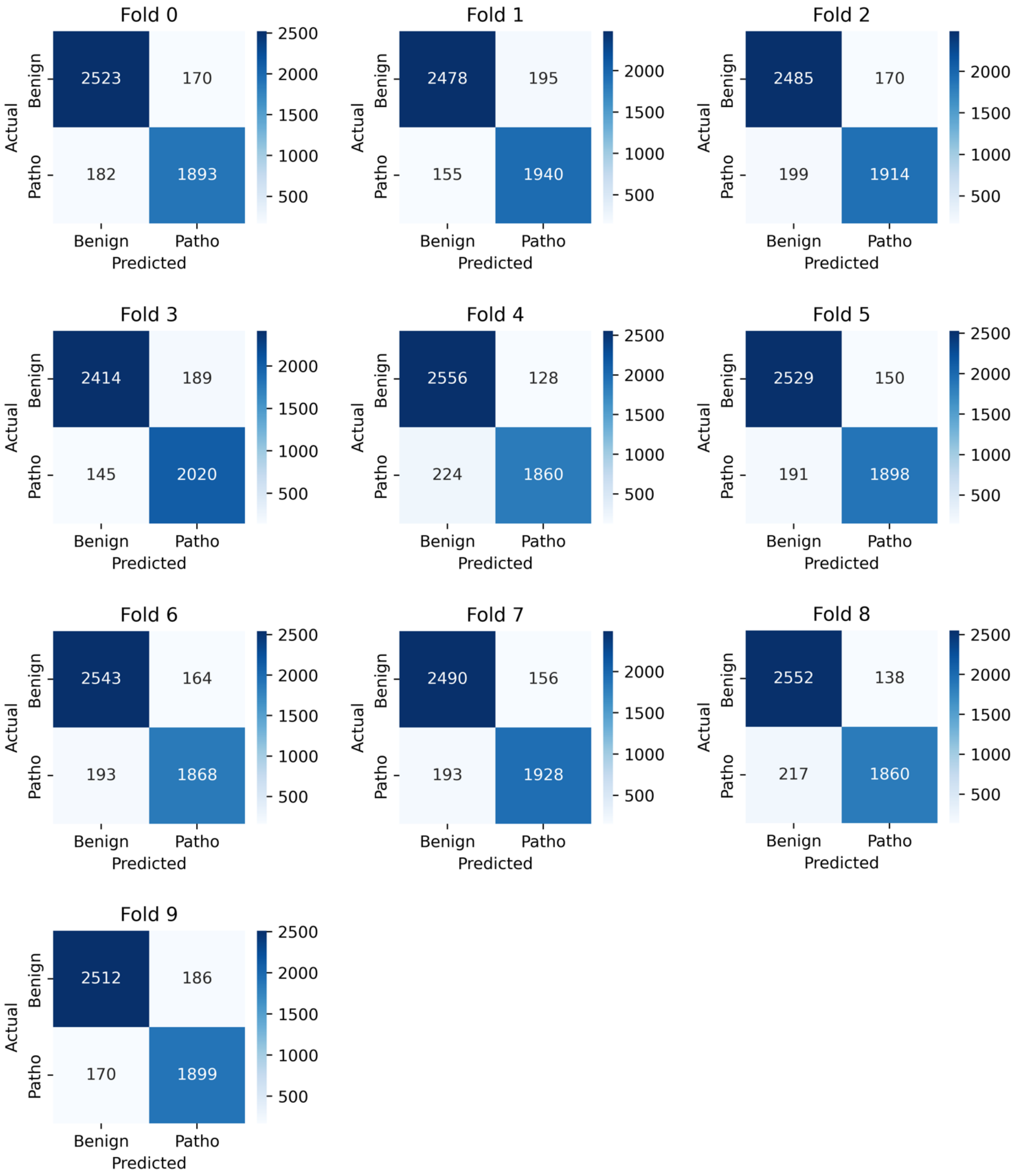


**Supplementary Figure 5.** Confusion matrix of Memo-Patho's 10-fold cross-validation results on the Mix dataset.

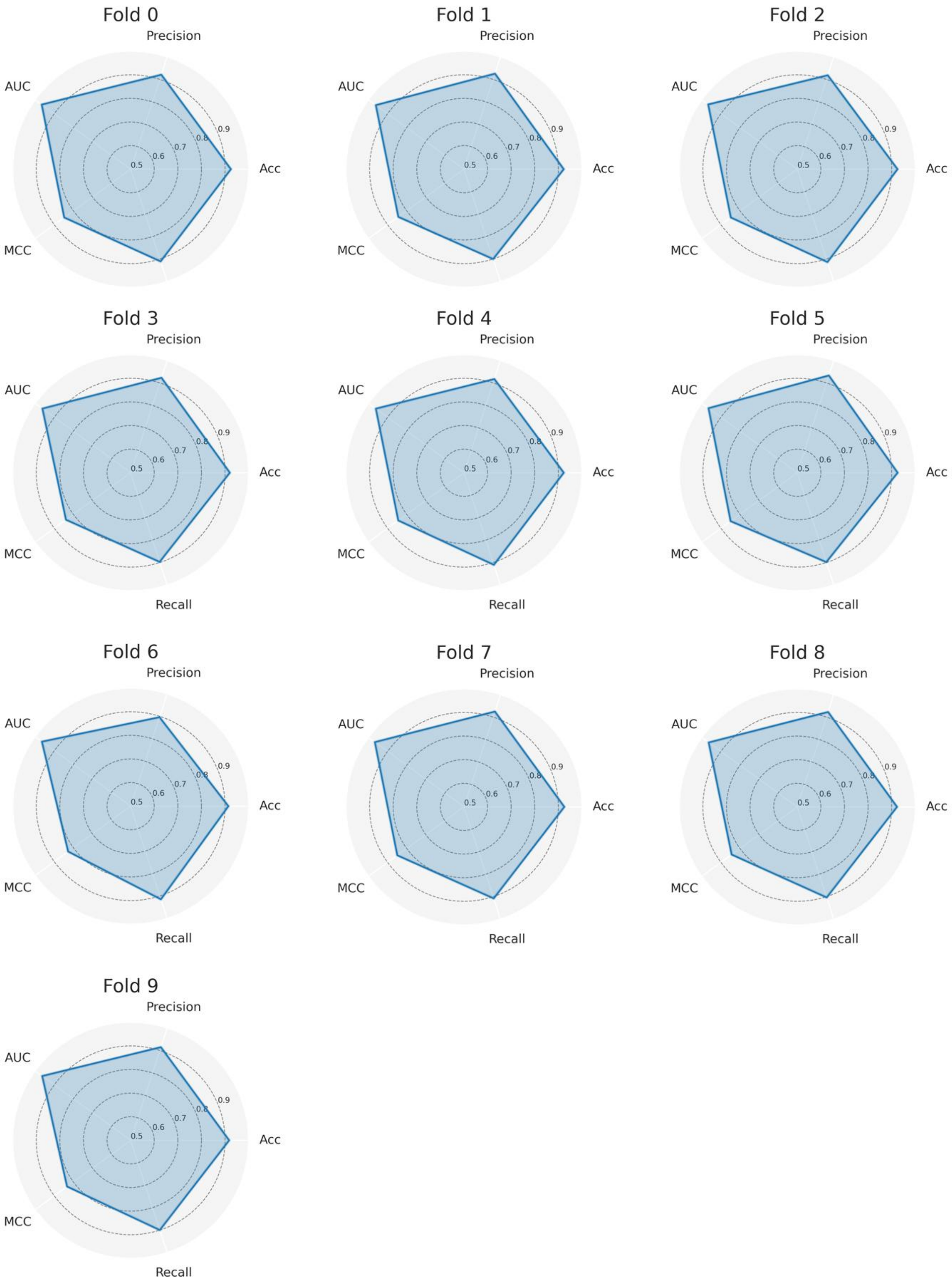


**Supplementary Figure 6.** Radar chart of Memo-Patho's 10-fold cross-validation results on the Mix dataset.

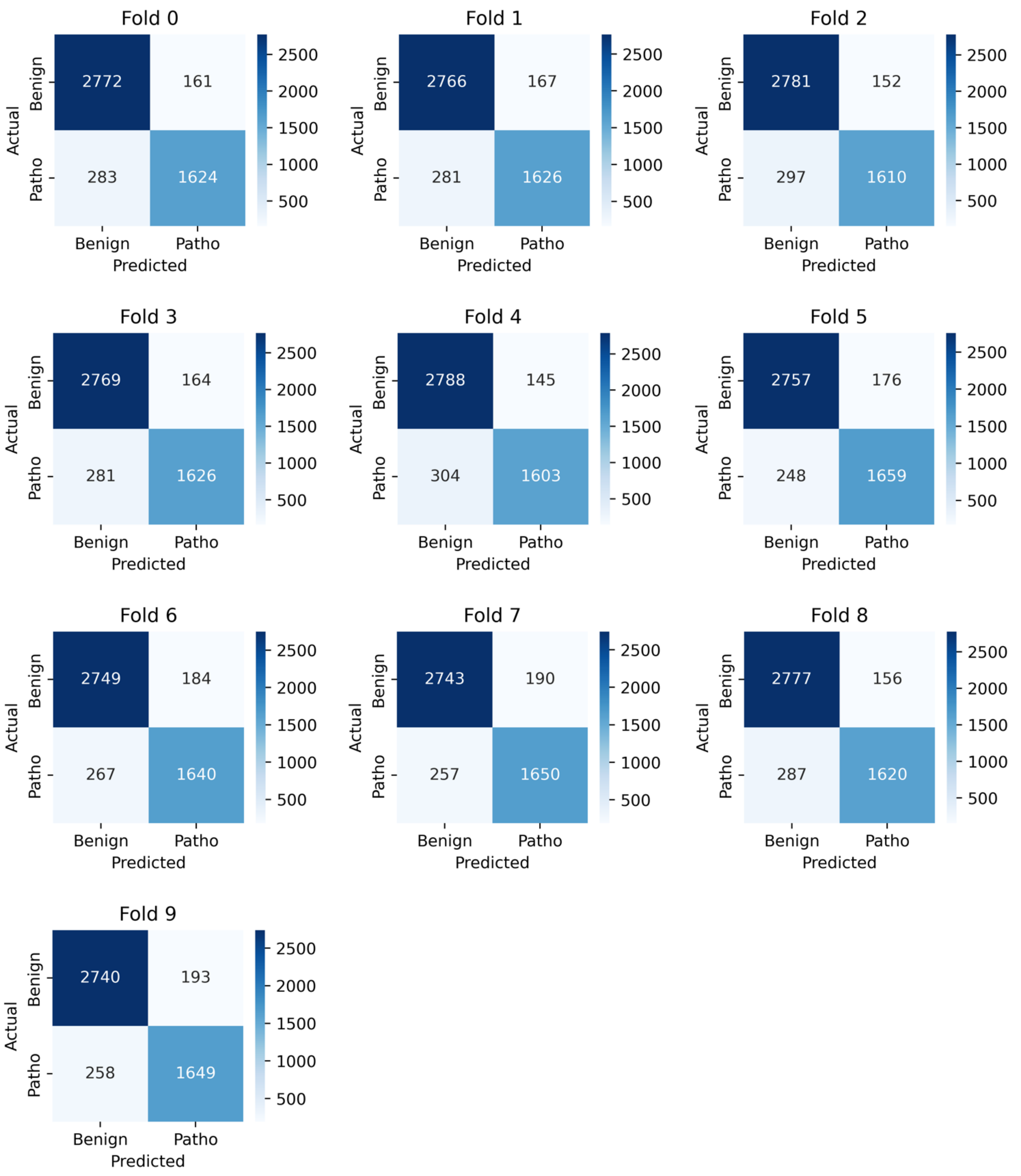


**Supplementary Figure 7.** Confusion matrix of Memo-Patho's 10-fold cross-validation results on the Ind dataset.

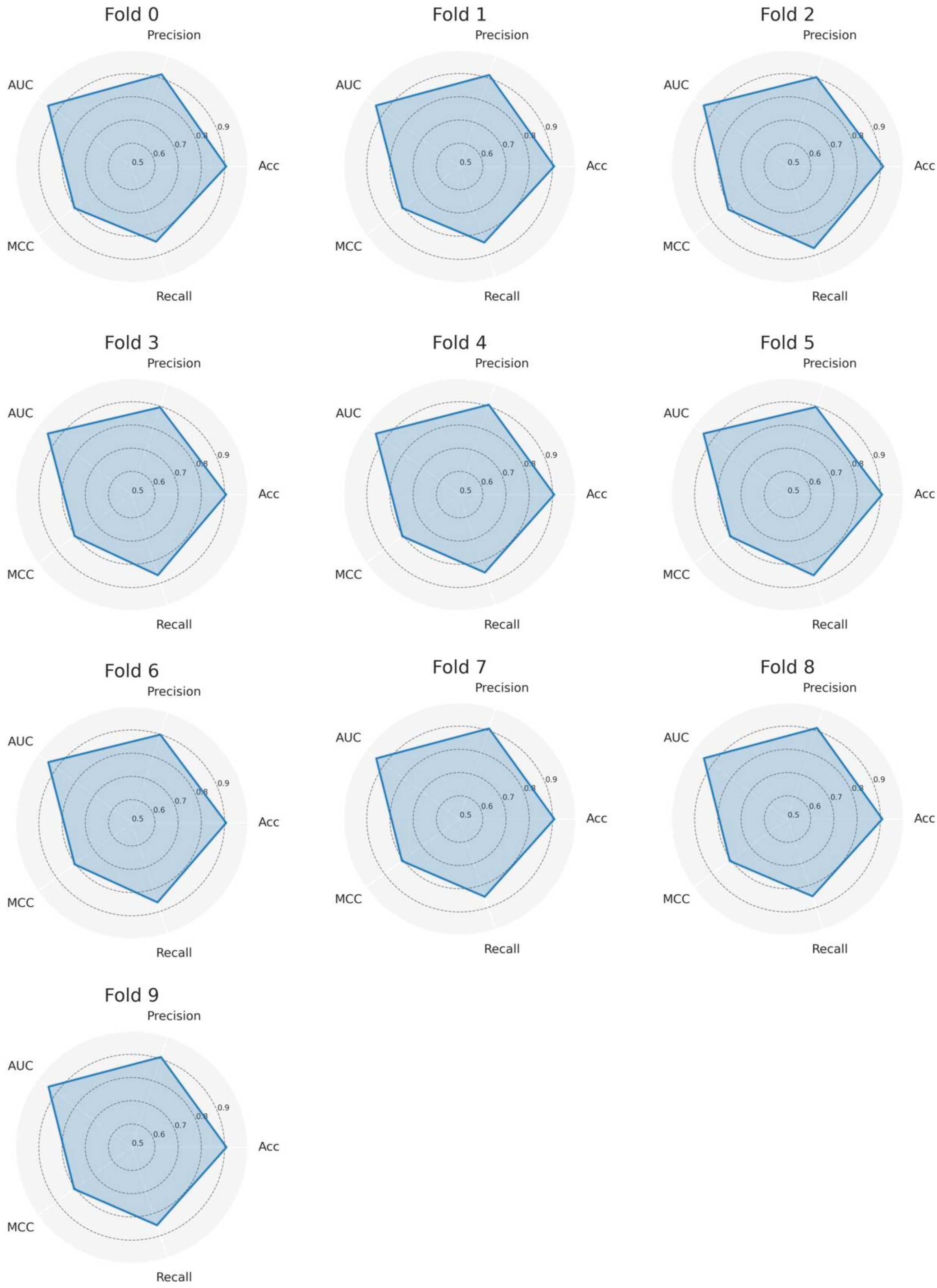


**Supplementary Figure 8.** Radar Chart of Memo-Patho's 10-fold cross-validation results on the Ind dataset.

**Supplementary Note**

**PLMs embedding generation**

1. ESM2: We employed the ESM2 protein language model in this study. The model setup and deployment strictly followed the procedures outlined in the official Facebook Research ESM GitHub repository, accessible at https://github.com/facebookresearch/esm. Specifically, the pre-trained weights corresponding to the esm2_t36_3B_UR50D version were used. These weights were obtained directly from the official distribution point: https://dl.fbaipublicfiles.com/fair-esm/models/esm2_t36_3B_UR50D.pt. This model architecture comprises 36 transformer layers and contains approximately 3 billion trainable parameters.

2. ProtT5: The ProtT5 model implementation was based on the guidelines and code provided in the ProtTrans GitHub repository (https://github.com/agemagician/ProtTrans). We utilized the ProtT5-XL-UniRef50 variant, leveraging its pre-trained weights. These weights are publicly available via the Hugging Face model repository, located at https://huggingface.co/Rostlab/prot_t5_xl_uniref50/tree/main.

3. Justification for Model Selection: The selection of the esm2_t36_3B_UR50D and ProtT5--.0XL-UniRef50 models was guided by the objective to maximize predictive performance while operating within the limitations of our computational infrastructure. These models represented the most powerful options compatible with our hardware constraints at the time of the study.

**Calculation methods for evaluation metrics**

$$Accuracy = \frac{TP + TN}{TP + FN + FP + TN}$$

$$Recall = \frac{TP}{TP + FN}$$

$$Precision = \frac{TP}{TP + FP}$$

$$F1 - score = 2 \times \frac{Recall \times Precision}{Recall + Precision}$$

$$MCC = \frac{TP \times TN - FP \times FN}{\sqrt{(TP + FP)(TP + FN)(TN + FP)(TN + FN)}}$$

***Supplementary Table 1.*** *Performance evaluation between baseline models on Mix dataset*

| Model | Accuracy | Precision | Recall | F1 | MCC |
|---|---|---|---|---|---|
| Memo-Patho | 0.9317 | 0.9318 | 0.9146 | 0.9231 | 0.8618 |
| PON-P3 | 0.8128 | 0.7678 | 0.8144 | 0.7904 | 0.6225 |
| ESNPs&GO | 0.8769 | 0.9373 | 0.8464 | 0.8606 | 0.7507 |
| PROVEAN | 0.8001 | 0.7370 | 0.8255 | 0.7787 | 0.6007 |
| TransEFVP | 0.8327 | 0.8377 | 0.7778 | 0.8067 | 0.6611 |
| AlphaMissense | 0.8432 | 0.9207 | 0.7567 | 0.8057 | 0.6792 |
| MutPred2 | 0.8068 | 0.7656 | 0.8194 | 0.7916 | 0.6132 |
| PredMutHTP | 0.7695 | 0.6757 | 0.8868 | 0.767 | 0.5669 |
| PON-P3* | 0.6199 | 0.5524 | 0.6486 | 0.5967 | 0.2444 |
| AlphaMissense* | 0.8046 | 0.7973 | 0.7308 | 0.7626 | 0.5987 |

‘*’ indicates we treat ‘ambiguous’ sample as wrong.

***Supplementary Table 2.*** *Performance evaluation between baseline models on Ind dataset*

| Model | Accuracy | Precision | Recall | F1 | MCC |
|---|---|---|---|---|---|
| Memo-Patho | 0.9149 | 0.9118 | 0.8679 | 0.8893 | 0.8209 |
| PON-P3 | 0.8277 | 0.7374 | 0.8552 | 0.792 | 0.6515 |
| ESNPs&GO | 0.8932 | 0.8655 | 0.8459 | 0.8556 | 0.771 |
| PROVEAN | 0.8017 | 0.698 | 0.8167 | 0.7527 | 0.5939 |
| TransEFVP | 0.8117 | 0.8272 | 0.6601 | 0.7343 | 0.6004 |
| AlphaMissense | 0.8666 | 0.8692 | 0.7700 | 0.8166 | 0.7156 |
| PredMutHTP | 0.7716 | 0.6517 | 0.8729 | 0.7462 | 0.5658 |
| PON-P3* | 0.5999 | 0.7678 | 0.8144 | 0.7904 | 0.6225 |
| AlphaMissense* | 0.8205 | 0.7840 | 0.7378 | 0.7602 | 0.6177 |

‘*’ indicates we treat ‘ambiguous’ sample as wrong.

***Supplementary Table 3.*** *Performance evaluation between baseline models on novel dataset*

| **Model** | **Accuracy** | **Precision** | **Recall** | **F1** | **MCC** |
|---|---|---|---|---|---|
| Memo-Patho | 0.836 | 0.875 | 0.849 | 0.862 | 0.662 |
| ESNPs&GO | 0.764 | 0.727 | 0.970 | 0.831 | 0.520 |
| TransEFVP | 0.782 | 0.839 | 0.788 | 0.813 | 0.554 |
| AlphaMissense | 0.655 | 0.750 | 0.636 | 0.689 | 0.312 |
| PredMutHTP | 0.727 | 0.696 | 0.968 | 0.810 | 0.441 |
